# Mitochondria directly sense osmotic stress to trigger rapid metabolic remodeling via PDH regulation

**DOI:** 10.1101/2022.06.29.498104

**Authors:** Takeshi Ikizawa, Kazutaka Ikeda, Makoto Arita, Shojiro Kitajima, Tomoyoshi Soga, Hidenori Ichijo, Isao Naguro

**Author notes:** Correspondence (H.I.), (I.N.).

## Abstract

A high-salt diet significantly impacts various diseases, including cancer and immune diseases. Recent studies suggest that the high-salt/hyperosmotic environment in the body may alter the chronic properties of cancer and immune cells in the disease context. However, little is known about the acute metabolic changes in hyperosmotic stress. Here, we found that hyperosmotic stress for a few minutes induces Warburg-like metabolic remodeling in HeLa and Raw264.7 cells and suppresses fatty acid oxidation. Regarding Warburg-like remodeling, the PDH phosphorylation status was altered bidirectionally (high in hyperosmolarity and low in hypoosmolarity) to osmotic stress in isolated mitochondria, suggesting that mitochondria themselves have an acute osmo-sensing mechanism. Warburg-like remodeling is required for HeLa cells to maintain ATP levels and survive under hyperosmotic conditions. Our findings suggest that cells exhibit acute metabolic remodeling under osmotic stress via the regulation of PDH phosphorylation by direct osmosensing within mitochondria.

## Introduction

In modern society, people are increasingly consuming high-salt diets. High-salt diets are known to have negative effects on human health and exacerbate various diseases, such as hypertension, cardiovascular disease, autoimmune disease, and cancer (Mu *et al*, 2011; Gaddy *et al*, 2013; Kleinewietfeld *et al*, 2013; Mozaffarian *et al*, 2014). In particular, many studies have shown a relationship between high-salt diets and cancer, such as the increased risk of gastric cancer and accelerated progression and metastasis of breast cancer by the activation of the MAPK/ERK pathway in cancer cells (Tsugane and Sasazuki, 2007; Chen *et al*, 2020). In contrast, however, there are reports suggesting that feeding a high-salt diet to mice harboring tumors enhanced antitumor immunity and inhibited tumor progression by suppressing the activity of myeloid-derived suppressor cells (MDSCs) (Willebrand *et al*, 2019; He *et al*, 2020). Therefore, the effect of high-salt diets on cancer still remains unclear, and research is necessary to determine how high-salt conditions affect cancer cells and/or surrounding cells and what molecular mechanisms drive these effects.

It has been previously reported that the concentration of sodium ions is increased in tumor tissue (Leslie *et al*, 2019). For example, there is a higher accumulation of sodium in human breast cancer tissue than in other areas of the breast tissue (Ouwerkerk *et al*, 2007). In addition, an elevated sodium concentration was observed in malignant brain tumors (Ouwerkerk *et al*, 2003). Even though the precise mechanisms of sodium ion accumulation in tumor tissues are not clear, it is suggested that changes in the expression and/or activity of membrane channels and transporters are involved (Leslie *et al*, 2019). Under chronic hyperosmotic conditions, it has been reported that aerobic glycolysis in cancer cells is enhanced (Stubblefield and Mueller, 1960). In a breast cancer cell line, elevated sodium increases the expression of enzymes in glycolysis, such as hexokinase (HK) and lactate dehydrogenase (LDH), resulting in increased lactate production (Amara, Zheng and Tiriveedhi, 2016). However, little is known about the acute alteration in glucose and fatty acid metabolism and its underlying mechanisms upon hyperosmotic stress in cancer cells.

Myeloid-derived cells, such as macrophages, infiltrate the tumor microenvironment, and they are known to exhibit plasticity in response to environmental cues. In recent years, it has been suggested that in addition to the kidney, a high-sodium environment is observed in various sites of our body and regulates the immune system, implying that a hyperosmotic environment regulates immunity (Schatz *et al*, 2017; Müller *et al*, 2019). It has been reported that hyperosmotic conditions promote the differentiation of monocytes into proinflammatory M1 macrophages (Wilck *et al*, 2019; Geisberger *et al*, 2021). Furthermore, previous reports suggested in mouse models that a high-salt diet leads to the accumulation of sodium in the tumor and enhances antitumor immunity (Willebrand *et al*, 2019; He *et al*, 2020). Immunometabolism has revealed that the metabolic state of immune cells critically affects their polarization and function, including that of tumor-associated macrophages (TAMs) (Russell, Huang and VanderVen, 2019; Pan *et al*, 2020; Liu *et al*, 2021). Thus, it is an interesting issue whether the hyperosmotic microenvironment might affect the metabolism of infiltrating immune cells in the tumor.

In the present study, we demonstrate that glucose metabolic remodeling from oxidative phosphorylation (OXPHOS) to aerobic glycolysis (a Warburg-like effect) is rapidly triggered upon hyperosmotic stress both in HeLa cells (a cancer cell line) and in RAW264.7 cells (a mouse monocyte cell line). Moreover, we discovered that fatty acid oxidation (FAO) is acutely suppressed, and acyl-carnitine accumulates under hyperosmotic conditions. We found that phosphorylation of PDH is upregulated and downregulated quickly upon hyperosmotic and hypoosmotic stress, respectively, both in cells and in isolated mitochondria. Our results suggest that Warburg-like metabolic remodeling is induced through the rapid regulation of PDK-dependent PDH phosphorylation upon hyperosmotic stress, which is directly sensed by mitochondria. Collectively, our results provide new insight into rapid metabolic remodeling upon osmotic stress, which might enable us to regulate cellular metabolism intentionally and rapidly by changing the environmental osmolarity.

## Results and Discussion

### Warburg-like metabolic remodeling is rapidly induced upon hyperosmotic stress

To investigate the rapid change in cellular metabolism upon hyperosmotic stress, we measured the oxygen consumption rate (OCR) and extracellular acidification rate (ECAR), which indicate the speed of mitochondrial respiration and aerobic glycolysis, respectively. We found that OCR markedly decreased upon hyperosmotic stress (500 mOsm) by adding 100 mM sodium chloride to HeLa cells and RAW264.7 cells (Fig 1A). It was a prompt response that could be almost saturated within 10 minutes (a measurement interval) after stimulation. Oligomycin-sensitive basal OCR (ΔOCR_Oligo_), which is coupled to mitochondrial ATP production, was significantly suppressed upon hyperosmotic stress (Fig 1B). Moreover, maximal respiration, which can be derived from an increase upon FCCP treatment, was also significantly suppressed upon hyperosmotic stress (Fig 1C). We also observed that ECAR rapidly increased upon hyperosmotic stress in HeLa cells (Fig 1D, E), as previously reported (Epstein *et al*, 2014). As ECAR evaluates protons mainly derived from lactate, the conversion of pyruvate to lactate seemed to be activated upon hyperosmotic stress, similar to the Warburg effect (Palsson-mcdermott, 2013). A decrease in OCR and an increase in ECAR (a Warburg-like effect) were also observed by adding 200 mM sorbitol (500 mOsm) as well as sodium chloride stimulation (Fig EV1A-E). These results suggest that the observed Warburg-like effect is triggered by hyperosmolarity but not by a high concentration of sodium chloride itself.

**Figure 1.**
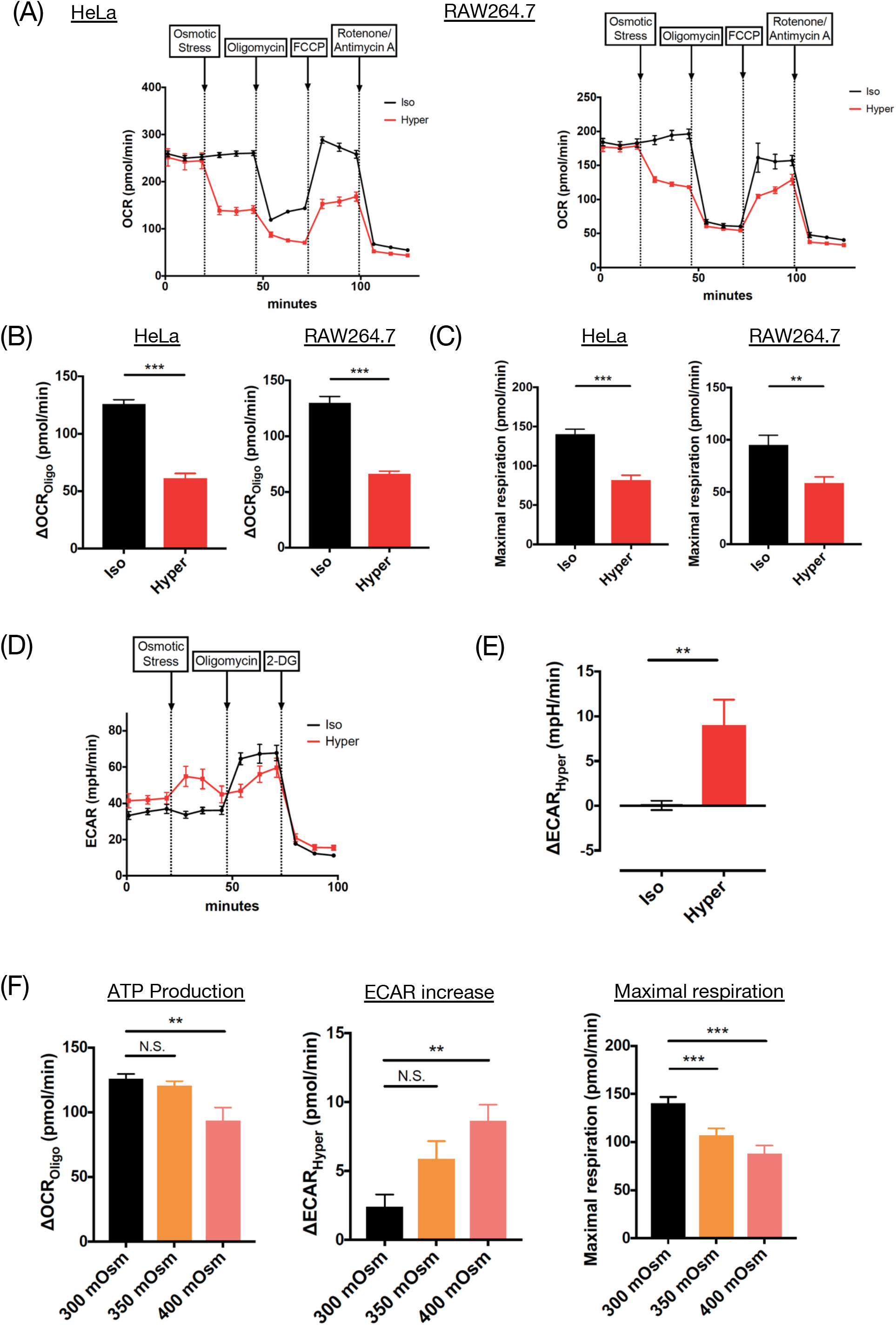
Acute Warburg-like metabolic remodeling is induced upon hyperosmotic stress. (A) OCR of HeLa and RAW264.7 cells was measured with sequential treatment with osmotic stress, oligomycin (ATP synthase inhibitor), FCCP (uncoupler), and rotenone (complex I inhibitor)/antimycin A (complex ? inhibitor) using an extracellular flux analyzer. A representative result from three independent experiments of each cell line is shown. The trace was the average of 3-10 independent wells for each condition. (B) Oligomycin-dependent OCR (ΔOCR_Oligo_) upon osmotic stress extracted from (A), which was calculated by subtraction between the average of three sequential measurements before and after oligomycin. (C) Maximal respiration upon osmotic stress extracted from (A), which was calculated by subtraction between the average of three sequential measurements before and after FCCP. (D) ECAR of HeLa cells was measured with sequential treatment with osmotic stress, oligomycin, and 2-DG (hexokinase inhibitor) using an extracellular flux analyzer. A representative result from five independent experiments is shown. The trace was the average of 3-5 independent wells for each condition. (E) Increase in ECAR by hyperosmotic stress (ΔECAR_Hyper_) extracted from (D), which was calculated by subtraction between the average of three sequential measurements before and after osmotic stress. (F) ΔOCR_Oligo_, ΔECAR_Hyper_ and maximal respiration of HeLa cells under 300, 350 and 400 mOsm. Three independent experiments were performed in which 3-10 independent wells were measured for each condition. Isoosmotic stress: 300 mOsm, Hyperosmotic stress: 500 mOsm. Data are represented as the mean ± SEM. ** p < 0.01, *** p < 0.001. Unpaired two-tailed Student’s t-test for (B), (C), (E). One-way ANOVA followed by Dunnett’s multiple comparisons test for (F).

Then, we examined these metabolic changes under milder hyperosmotic stress (Fig 1F). Treatment with 50 mM sodium chloride (400 mOsm) significantly suppressed ΔOCR_Oligo_ and increased ΔECAR_Hyper_. The maximal respiration was also significantly suppressed. In the case of 25 mM sodium chloride (350 mOsm), although the suppression of basal OCR and enhancement of ECAR were marginal, maximal respiration was significantly suppressed. Considering that the hyperosmotic environment observed in the human body (except the kidney, exhibiting extremely high osmolarity) reaches up to 400 mOsm (Go *et al*, 2004; Ouwerkerk *et al*, 2007; Wiig *et al*, 2013; Jantsch *et al*, 2015; Hara *et al*, 2017) the Warburg-like effect observed upon hyperosmotic stress is likely to occur under physiological conditions of various tissues.

As changes in OCR and ECAR were observed dose-dependently by hyperosmotic stress, we further investigated the responses to hypoosmotic stress. Interestingly, OCR significantly increased upon hypoosmotic stress (225 mOsm) (Fig EV1F, G). Additionally, OCR suppression by hyperosmolarity was gradually restored to the level of isoosmolarity when we added water to hyperosmotic medium to dilute it to isoosmotic conditions (300 mOsm). Taken together, these results suggest that OCR bidirectionally and flexibly responds to osmotic stress. Meanwhile, ECAR did not change upon hypoosmotic stress (Fig EV1H).

### Glucose metabolism shifts toward aerobic glycolysis upon hyperosmotic stress

To examine the contribution of glucose metabolites to the Warburg-like effect upon hyperosmotic stress, we conducted a metabolome analysis using a fully labeled ^13^C_6_-glucose tracer. Among the detected metabolites in glycolysis, metabolites toward phosphoenolpyruvate (PEP) were mostly replaced with the full-labeled form within 10 minutes in both isoosmotic and hyperosmotic conditions (Fig 2A). The amount of full-labeled fructose 1,6-bisphosphate (F1,6P) was significantly increased under hyperosmotic conditions compared with isoosmotic conditions at 30 and 60 minutes, although there was no change in the amount of full-labeled fructose 6-phosphate (F6P) (Fig 2A). These data suggest that phosphofructokinase (PFK) might be activated under hyperosmotic stress with a sufficient supply of F6P from glucose. Moreover, the total amount and full-labeled form of intermediate metabolites in glycolysis, dihydroxyacetone phosphate (DHAP), 2,3-diphosphoglycerate (2,3-DPG), 3-phosphoglycerate (3PG), and PEP, were also increased under hyperosmotic conditions compared with isoosmotic conditions (Fig 2A, EV2A). Finally, the end product of aerobic glycolysis, lactate, showed an increasing tendency in total amount and a significant increase in full-labeled form at 30 minutes in hyperosmotic conditions, although pyruvate did not significantly increase in total amount and full-labeled form, suggesting that the conversion of pyruvate to lactate was activated upon hyperosmotic stress. This is consistent with the increase in ECAR upon hyperosmotic stress (Fig 1D, E).

**Figure 2.**
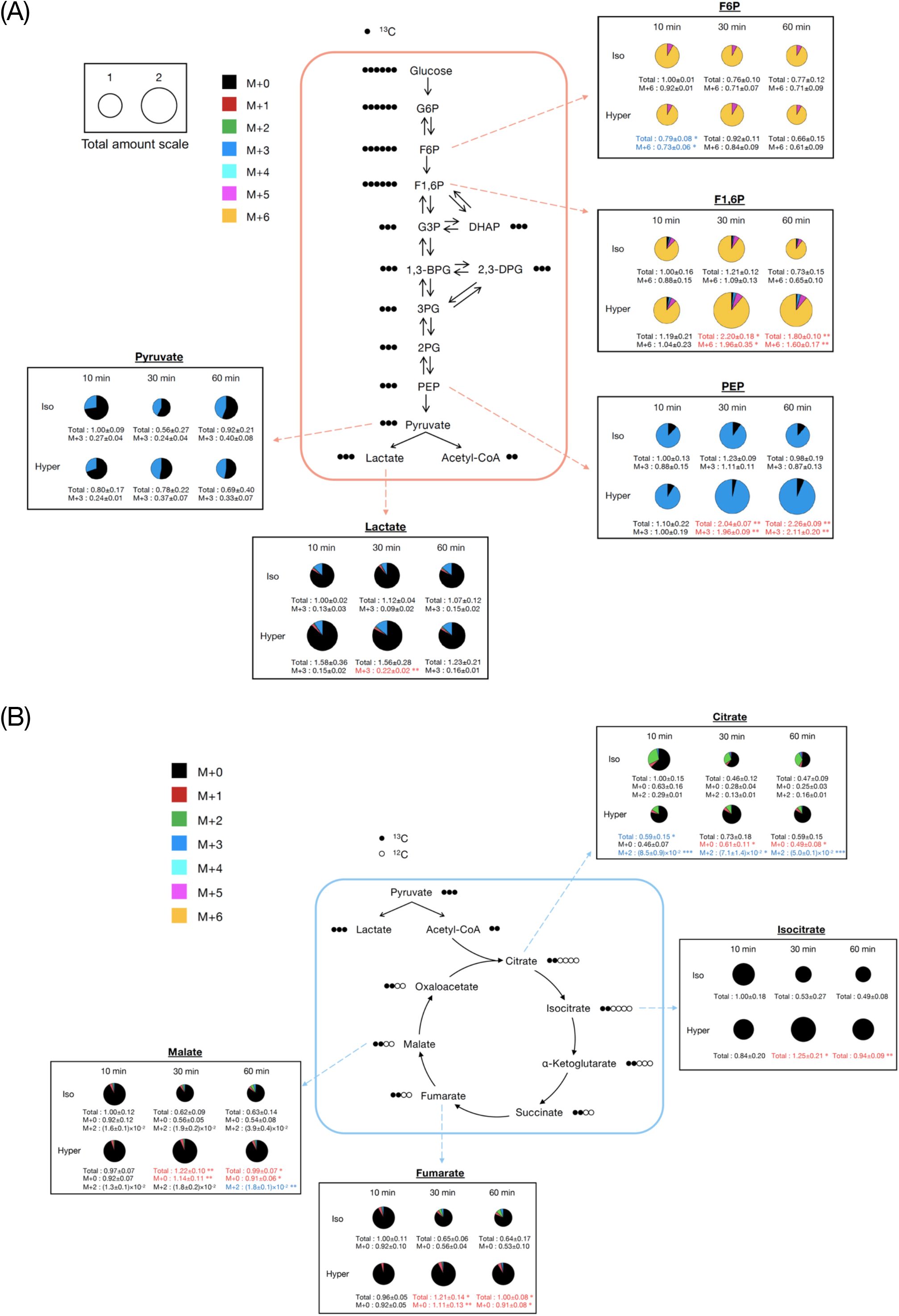
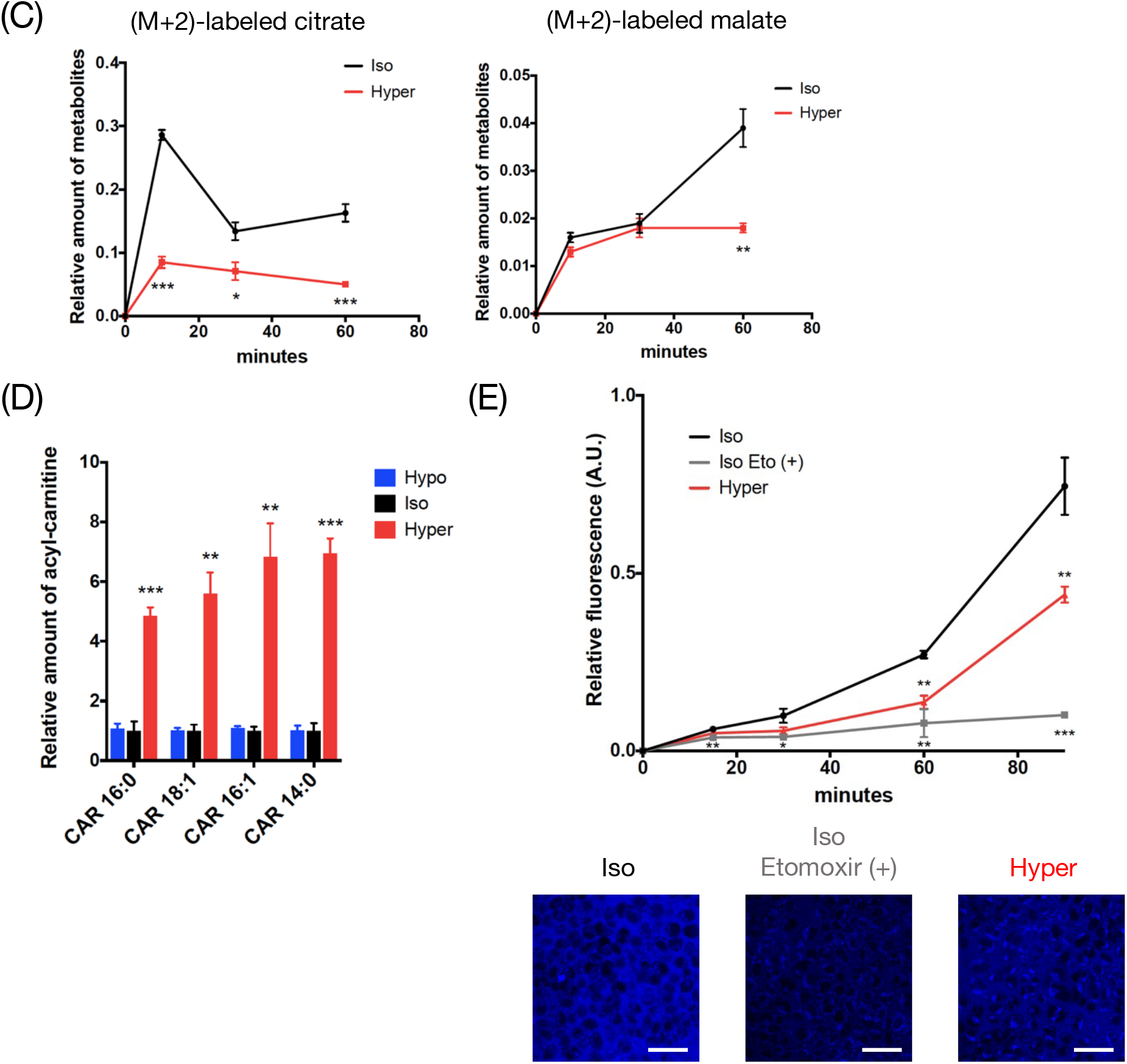
Aerobic glycolysis is activated and the supply of acetyl-CoA to the TCA cycle is reduced upon hyperosmotic stress. (A, B) Metabolome analysis of HeLa cells using ^13^C_6_-glucose tracer under isoosmotic and hyperosmotic conditions for 10, 30 and 60 min. For each metabolite, the relative amounts are normalized by that of the total amount at 10 min under isoosmotic conditions (that is, the total amount at 10 min is 1). The size of each circle depicts the relative amount of a metabolite in total (nonlabeled to full-labeled). The size scale of the circles and color codes of each metabolite are indicated in the upper left. The relative amounts of several metabolites labeled with a specific number of ^13^C are shown by text under each circle. Metabolites significantly increased and decreased under hyperosmotic conditions are shown in red and blue letters, respectively. Metabolites in glycolysis (F6P, F1,6P, PEP, pyruvate, and lactate) are shown in (A), and metabolites in the TCA cycle (citrate, isocitrate, fumarate, and malate) are shown in (B). (C) Line graphs of the relative amount of (M+2)-labeled citrate and malate extracted from (B). (D) Relative amount of acyl-carnitines (composed of 16:0, 18:1,16:1 and 14:0 fatty acids) detected by nontargeted lipidome analysis. Samples were harvested from HeLa cells after 15 min of osmotic stress. The amount of each acyl-carnitine (CAR n:n) is normalized to that in isoosmotic conditions. (E) Fluorescence intensity of FAOBlue in HeLa cells at 15, 30, 60 and 90 min after osmotic stress. Three independent experiments were performed in triplicate from independent wells. The upper line graph shows a representative result with the mean ± SEM of triplicate experiments. The lower panels show FAOBlue fluorescence images of HeLa cells at 90 min after osmotic stress. Etomoxir (40 µM) was pretreated before osmotic stress for 30 min in the experiment measuring fluorescence intensity and for 60 min in the live-cell imaging experiment. Scale bar: 50 µm. Hypoosmotic stress: 200 mOsm, Isoosmotic stress: 300 mOsm, Hyperosmotic stress: 500 mOsm. Data are represented as the mean ± SEM. N=3 (three samples for each condition) except (E). * p < 0.05, ** p < 0.01, *** p < 0.001. Unpaired two-tailed Student’s t-test at the same time points for (A)-(C). One-way ANOVA followed by Dunnett’s multiple comparisons test for (D), (E).

Regarding the metabolites in the TCA cycle, the amount of (M+2)-labeled citrate (containing two ^13^C), which is mainly derived from acetyl-CoA through glycolysis, was significantly suppressed upon hyperosmotic stress (Fig 2B, C). The amount of (M+2)-labeled malate was also suppressed under hyperosmotic conditions at 60 minutes. Although we could not precisely measure acetyl-CoA in the metabolome analysis, defects in ^13^C incorporation into TCA cycle metabolites suggest that the supply of glucose-derived acetyl-CoA to the TCA cycle might be reduced under hyperosmotic conditions.

On the other hand, nonlabeled (M+0) citrate, isocitrate, fumarate and malate significantly accumulated at 30 and 60 min under hyperosmotic stress compared with isoosmotic conditions (Fig 2B). This result may reflect the previous observation that OXPHOS is directly impaired by hyperosmotic stress (Geisberger *et al*, 2021), resulting in a decrease in the rate of the TCA cycle. On the other hand, it is also possible that resources of acetyl-CoA other than glucose might expand in hyperosmotic conditions to increase nonlabeled metabolites in the TCA cycle. As fatty acids are another main source of acetyl-CoA (Kishton and Rathmell, 2017), we comprehensively examined lipid dynamics in osmotic conditions by a nontargeted lipidome analysis (Tsugawa *et al*, 2020). We found that various species of acyl-carnitine, an intermediate metabolite of fatty acid metabolism in mitochondria (β-oxidation), markedly increased at 15 minutes of hyperosmotic stress (Fig 2D). There was no difference in the amount of any kind of triglycerides, which are sources of acyl-carnitine (Fig EV2B). Then, we quantified the fatty acid oxidation activity in HeLa cells upon osmotic stress using FAOBlue, the fluorescence of which reflects the β-oxidation activity in live cells (Uchinomiya *et al*, 2020). FAOBlue fluorescence intensity, which was abolished by etomoxir (an inhibitor of carnitine palmitoyl transferase 1 (CPT1)), was attenuated under hyperosmotic stress (Fig 2E). Furthermore, in culture medium with rich palmitic acid and no glucose, the maximal respiration (depending on fatty acid metabolism) was suppressed under hyperosmotic stress (Fig EV2C). Collectively, these results indicate that β-oxidation was suppressed by hyperosmotic stress, in parallel with the accumulation of acyl-carnitine. Therefore, it is conceivable that acetyl-CoA derived from fatty acids was also decreased under hyperosmotic conditions, suggesting that the accumulation of nonlabeled metabolites in the TCA cycle (seen in Fig 2B) was due to the reduction in consumption in the TCA cycle.

These results also suggest that, as well as glucose metabolic remodeling, the suppression of β-oxidation contributes to the decrease in OCR through the reduction of acetyl-CoA supply. On the other hand, it remains unclear whether the accumulation of acyl-carnitine is the result of the suppression of β-oxidation. The defect in *β*-oxidation would simply result in the accumulation of a substrate, acyl-carnitine. It is also conceivable that impairment of acyl-carnitine transport to the mitochondrial matrix caused by defects in carnitine acyl-carnitine translocase (CACT) and/or carnitine palmitoyl transferase 2 (CPT2) may abrogate β-oxidation. Thus, the molecular mechanism by which hyperosmotic stress increases acyl-carnitine in the cells and the physiological meaning of the inhibition of fatty acid metabolism have yet to be elucidated.

### PDH is suppressed through phosphorylation by PDK upon hyperosmotic stress, which is directly sensed by mitochondria

The decrease in OCR upon hyperosmotic stress was attributable to a reduction in acetyl-CoA supply (from glucose and fatty acids) and/or defects in the electron transport chain (Geisberger *et al*, 2021). First, we evaluated the contribution of acetyl-CoA supply from glucose and fatty acid metabolism to the decrease in OCR under normal culture conditions. Surprisingly, prior inhibition of glycolysis by 2-deoxy-glucose (2-DG) strongly abrogated the decrease in OCR upon hyperosmotic stress, suggesting that the alteration in glycolysis is the main reason for the acute decrease in OCR (Fig 3A, B). Considering that OCR was almost maintained even under hyperosmotic stress in 2-DG conditions, in contrast to a previous report regarding mononuclear phagocytes (Geisberger *et al*, 2021), a direct defect in the electron transport chain is not the main reason for the acute OCR decrease upon hyperosmotic stress in the present context. In contrast, prior inhibition of β-oxidation by etomoxir treatment scarcely abolished the decrease in OCR induced by hyperosmotic stress (Fig 3C, D). These results suggest that under normal culture conditions, suppression of acetyl-CoA supply via glycolysis plays a pivotal role in acute OCR suppression upon hyperosmotic stress.

**Figure 3.**
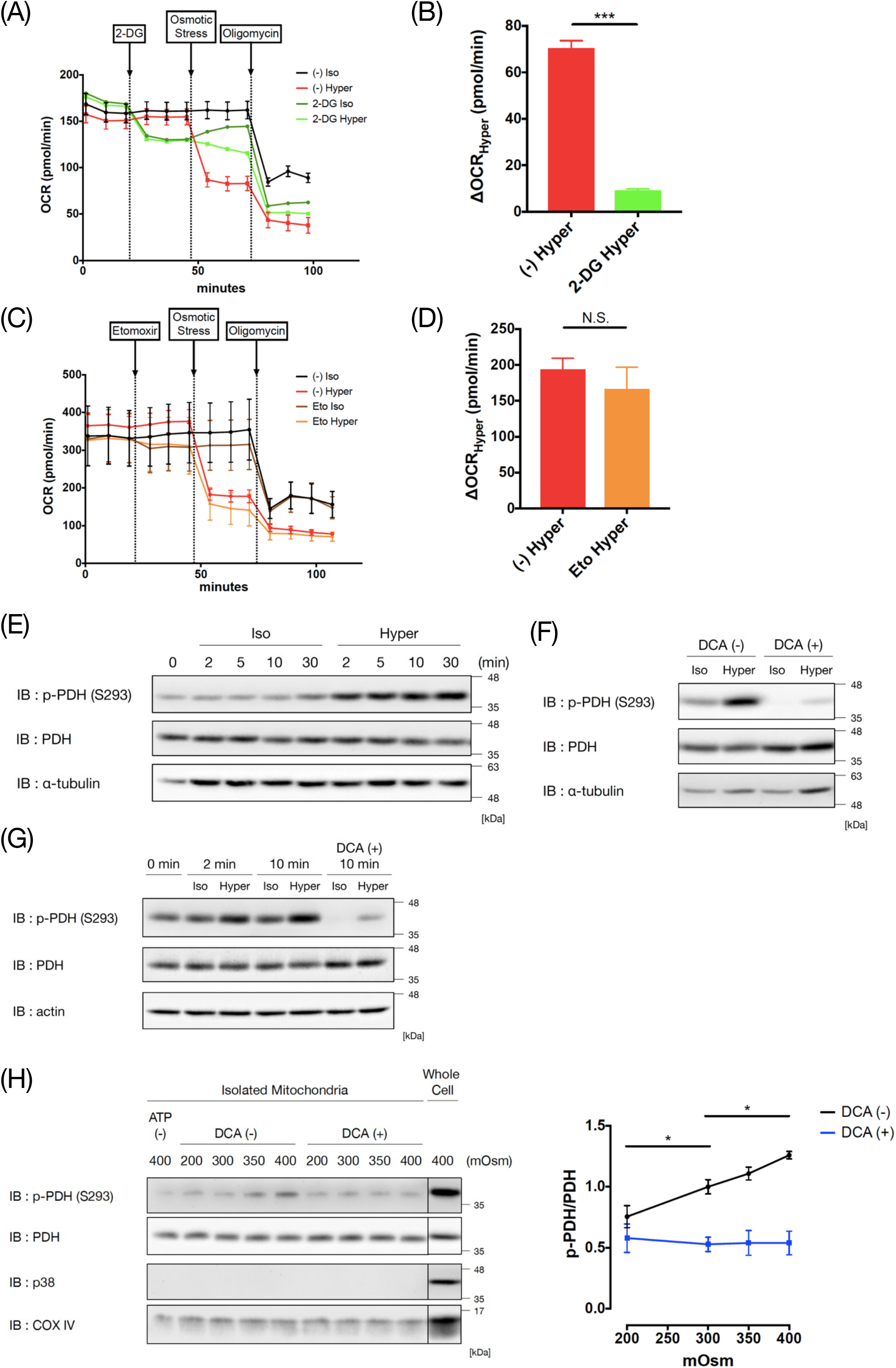
PDK-dependent PDH phosphorylation is induced in mitochondria upon hyperosmotic stress. (A), (C) OCR of HeLa cells with sequential treatment with 2-DG (30 mM) (A) or etomoxir (20 µM) (C), osmotic stress and oligomycin. A representative result from three independent experiments is shown. The trace was the average of 3-7 (A) or 3-10 (C) independent wells for each condition. (B), (D) Decrease in OCR by hyperosmotic stress (ΔOCR_Hyper_) extracted from (A) or (C), respectively, which was calculated by subtraction between the average of three sequential measurements before and after osmotic stress. (E) Immunoblot analysis of PDH phosphorylation after 2, 5, 10 and 30 min of osmotic stress in HeLa cells. (F) Immunoblot analysis of PDH phosphorylation after 10 min of osmotic stress with and without pretreatment with DCA (25 mM) for 60 min in HeLa cells. (G) Immunoblot analysis of PDH phosphorylation after 2 and 10 min of osmotic stress in RAW264.7 cells. DCA (25 mM) was pretreated for 60 min. (H) Immunoblot analysis of PDH phosphorylation in isolated mitochondria from HeLa cells. Isolated mitochondria were directly exposed to 10 min of osmotic stress (200, 300, 350 and 400 mOsm). DCA (25 mM) was pretreated for 30 min. ATP (500 nM) was added at the same time as osmotic stimulation. Whole-cell samples of HeLa cells were prepared after 30 min of hyperosmotic stimulation (400 mOsm). The right graph shows the ratio of p-PDH to PDH quantified from the immunoblot. Three independent experiments were performed for quantification of the band intensities. Isoosmotic stress: 300 mOsm, Hyperosmotic stress: 500 mOsm. Data are represented as the mean ± SEM. * p < 0.05, *** p < 0.001. Unpaired two-tailed Student’s t-test for (B), (D). One-way ANOVA followed by Dunnett’s multiple comparisons test for (H).

Then, we hypothesized that pyruvate dehydrogenase (PDH), which converts pyruvate to acetyl-CoA, might be inactivated under hyperosmotic stress, similar to the general Warburg effect (Woolbright *et al*, 2019). PDH is inactivated in various conditions, such as hypoxia or nutrient starvation, by the phosphorylation of serine residues S232, S293 or S300 (Korotchkina and Patel, 2001). We found that the phosphorylation status of PDH(S293) was upregulated upon hyperosmotic stress (Fig 3E). The upregulation was observed as early as 2 minutes and was sustained until at least 30 minutes upon hyperosmotic stimulation. This result suggests that PDH inactivation may suppress the acetyl-CoA supply from pyruvate, resulting in the activation of lactate production. PDH(S293) phosphorylation was also upregulated upon hyperosmotic stress by adding sorbitol, confirming the importance of osmolarity itself (Fig EV3A).

Interestingly, corresponding to the OCR increase under hypoosmotic stress (Fig EV1E), PDH(S293) phosphorylation decreased upon hypoosmotic stress (Fig EV3B). The phosphorylation status of PDH(S293) was gradually altered in both directions in an osmolarity-dependent manner. Moreover, PDH(S293) phosphorylation was flexibly reversed after inversing osmotic stress (Fig EV3C). These results suggest that PDH(S293) phosphorylation is regulated bidirectionally and reversibly under osmotic stress.

PDH(S293) is phosphorylated by pyruvate dehydrogenase kinase (PDK) (Korotchkina and Patel, 2001). Pretreatment with dichloroacetate (DCA), a PDK inhibitor, strongly suppressed PDH(S293) phosphorylation upon hyperosmotic stress, suggesting that PDK is responsible for the phosphorylation of PDH(S293) under hyperosmotic stress (Fig 3F). Upregulation of PDH(S293) phosphorylation upon hyperosmotic stress and its dependence on PDK were also observed in RAW264.7 cells (Fig 3G). These results suggest that glucose metabolic remodeling is induced at least in part by the phosphorylation and inactivation of PDH by PDK with reversible and flexible plasticity.

It may be possible that the balance of glucose metabolism between OXPHOS and aerobic glycolysis can be intentionally and immediately controlled by changing the osmolarity, which is expected to provide a way to control the metabolic state of cancer cells and immune cells in cancer therapy. Moreover, activating aerobic glycolysis is one of the important remodeling pathways for macrophages to promote polarization into proinflammatory M1 macrophages (Russell, Huang and VanderVen, 2019). The tumor microenvironment is suggested to be hyperosmotic (Leslie *et al*, 2019) and that a high-salt diet is reported to result in further high-sodium conditions in tumors and inhibit tumor growth by enhanced tumor immunity (He *et al*, 2020). Therefore, it is possible that the infiltrating monocytes may enhance differentiation into M1 macrophages by activating aerobic glycolysis under such hyperosmotic conditions and may contribute to suppressing cancer progression by showing their antitumor activities.

Next, we examined the involvement of ERK, AMPK and Akt, which were previously reported to regulate PDK with their kinase activities (Wu *et al*, 2013; Cerniglia *et al*, 2016; Chae *et al*, 2016; Li *et al*, 2016; Cai *et al*, 2020). However, a MEK inhibitor (U0126) did not abrogate the phosphorylation of PDH(S293) upon hyperosmotic stress (Fig EV3D). The mechanism underlying the regulation of PDK by AMPK and Akt is unclear, and both positive and negative regulation have been reported (Wu *et al*, 2013; Cerniglia *et al*, 2016; Chae *et al*, 2016; Cai *et al*, 2020). AMPK activity monitored by S79 phosphorylation of ACC (Fig EV3E) and Akt activity monitored by S473 phosphorylation of Akt (Fig EV3F) were downregulated under hyperosmotic stress, implying that positive regulation of PDK by AMPK or Akt (Wu *et al*, 2013; Chae *et al*, 2016) could not be applicable under hyperosmotic conditions. AMPK activator (AICAR) did not suppress the phosphorylation of PDH(S293) upon hyperosmotic stress, although ACC phosphorylation was sustained by the treatment. This result suggests that negative regulation of PDK by AMPK does not contribute to the phosphorylation of PDH(S293) upon hyperosmotic stress (Fig EV3E). In addition, an Akt inhibitor (MK-2206) did not upregulate the phosphorylation of PDH(S293) under basal conditions, implying that Akt may not negatively regulate PDK in HeLa cells (Fig EV3F).

Considering that these cytoplasmic kinases upstream of PDK did not contribute to the regulation of PDH(S293) phosphorylation upon hyperosmotic stress and that both PDH and PDK are localized in mitochondria (Hitosugi *et al*, 2011), we next examined whether direct application of osmotic stress to mitochondria may alter the phosphorylation status of PDH(S293). We isolated mitochondria from HeLa cells and directly treated them with osmotic stress. The phosphorylation status of PDH was altered bidirectionally to osmotic stress and was inhibited by pretreatment with DCA (Fig 3H). These results suggest that mitochondria themselves can directly sense osmotic stress and change PDK activity. Furthermore, because no substrate for the TCA cycle was included in the reaction buffer, it is suggested that the alteration in PDK activity is independent of the secondary effect through changes in metabolites involved in glycolysis and the TCA cycle. It is possible that, similar to cell volume change under osmotic conditions (Hoffmann, Lambert and Pedersen, 2009), the properties of the mitochondrial membrane (morphology and tension) might change under osmotic stress and affect PDK activity. A previous report showed in yeast that the activity of target of rapamycin complex 2 (TORC2) is regulated by plasma membrane tension under osmotic stress (Riggi *et al*, 2018). A similar mechanism might govern PDK activity on the mitochondrial membrane. Further studies are required to elucidate the sensing mechanism of osmolarity in mitochondria to alter PDK activity.

### Warburg-like metabolic remodeling contributes to the maintenance of ATP amount and survival under hyperosmotic stress

To investigate the physiological meaning of rapid metabolic remodeling upon hyperosmotic stress, we focused on the cellular ATP level, assuming that the shift from OXPHOS to aerobic glycolysis may produce ATP in a faster manner (Liberti and Locasale, 2016). We used DCA and oxamate (LDH inhibitor) to suppress the Warburg-like metabolic remodeling observed under hyperosmotic stress. DCA suppresses the step of inhibitory phosphorylation of PDH, while oxamate suppresses the step of pyruvate to lactate conversion (Fig 4A). Both inhibitors inhibited Warburg-like metabolic remodeling under hyperosmotic stress. Pretreatment with DCA inhibited the decrease in OCR upon hyperosmotic stress, resulting in a higher OCR under hyperosmotic conditions (Fig 4B). The ECAR was decreased in the basal state with DCA treatment and remained lower than that of the control upon hyperosmotic stress (Fig 4C). Oxamate increased the basal OCR, and the OCR remained higher than that of the control upon hyperosmotic stress (Fig 4D). The increase in ECAR upon hyperosmotic stress was inhibited by pretreatment with oxamate, resulting in a lower ECAR under hyperosmotic conditions (Fig 4E). These results suggest that both inhibitors can be used as suppressors of Warburg-like remodeling (lower OCR and higher ECAR) upon hyperosmotic stress.

**Figure 4.**
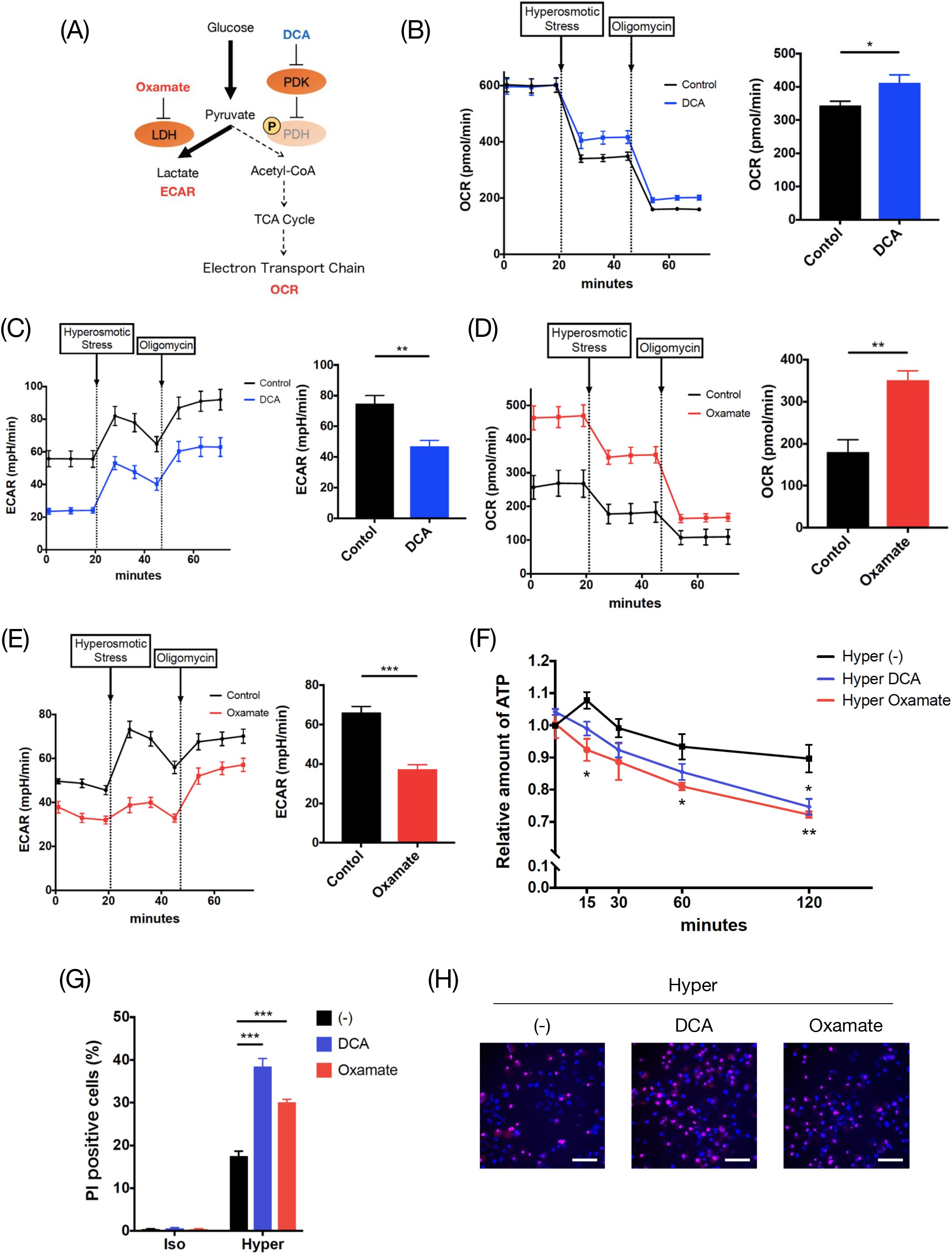
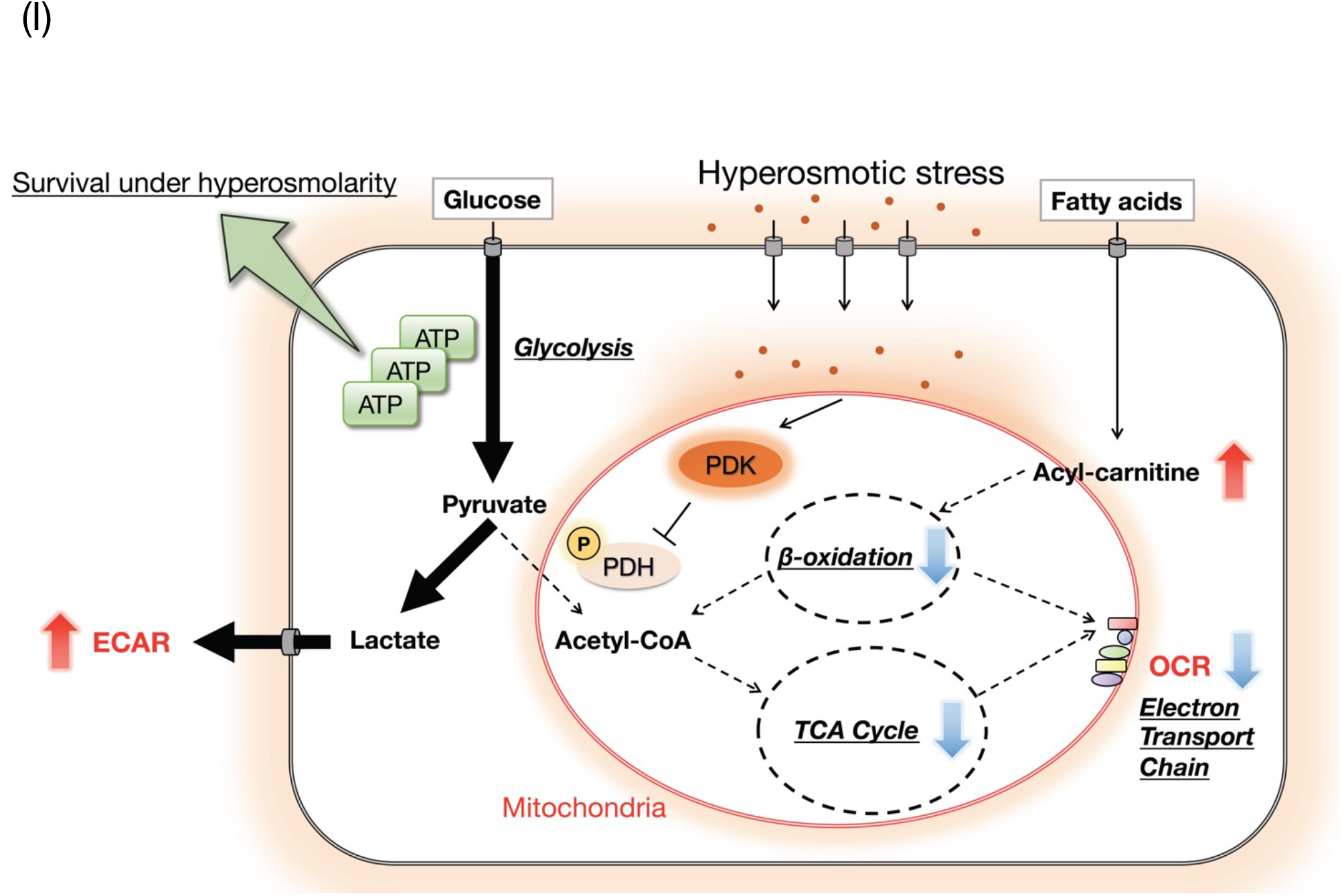
Warburg-like metabolic remodeling contributes to the maintenance of ATP levels and survival under hyperosmotic stress. (A) A schematic model of metabolic remodeling in hyperosmotic stress and molecular targets of DCA and oxamate. (B) OCR and (C) ECAR of HeLa cells upon hyperosmotic stress with pretreatment with NaCl (37.5 mM) or DCA (50 mM) for 30 min before the assay. NaCl was pretreated in the control to equalize the osmolarity with DCA treatment. A representative result from three independent experiments is shown. The trace was the average of 3-6 independent wells for each condition. Bar graphs show OCR (B) and ECAR (C) upon hyperosmotic stress, which was calculated by averaging the three sequential measurements upon hyperosmotic stress. (D) OCR and (E) ECAR of HeLa cells upon hyperosmotic stress with pretreatment with NaCl (37.5 mM) or oxamate (50 mM) for 30 min before the assay. NaCl was pretreated in the control to equalize the osmolarity with oxamate treatment. A representative result from three independent experiments is shown. The trace was the average of 3-6 independent wells for each condition. Bar graphs show OCR (D) and ECAR (E) upon hyperosmotic stress, which was calculated by averaging the three sequential measurements upon hyperosmotic stress. (F) The relative amount of ATP in HeLa cells was measured upon hyperosmotic stress with pretreatment with DCA (25 mM) or oxamate (25 mM) for 60 min using a plate reader. The timepoint of osmotic stimulation was defined as 0 min. Three independent experiments were performed and averaged. (G), (H) Percentage of PI-positive HeLa cells after 24 hr of osmotic stress with pretreatment with DCA (25 mM) or oxamate (25 mM) for 60 min measured by an image analyzer. Three independent experiments were performed and averaged in (G). Representative images of PI-stained HeLa cells under hyperosmotic conditions are shown in (H). Scale bar: 100 µm. (I) A schematic model of cellular metabolic changes upon hyperosmotic stress. When cells are subjected to hyperosmotic stress, the cytosol becomes hyperosmotic because of water efflux, resulting in an osmotic gradient between the cytosol and mitochondria. The osmotic change is directly sensed by the mitochondria and activates PDK, which phosphorylates PDH and suppresses its activity. This causes a shift in glucose metabolism from OXPHOS to aerobic glycolysis, resulting in a decrease in OCR and an increase in ECAR. Activation of aerobic glycolysis produces ATP quickly, which may contribute to cell survival under hyperosmotic conditions. Furthermore, fatty acid metabolism in mitochondria (*β*-oxidation) is suppressed, and acyl-carnitine accumulates upon hyperosmotic stress, which may be involved in the decrease in OCR. Isoosmotic stress: 300 mOsm, Hyperosmotic stress: 500 mOsm. Data are represented as the mean ± SEM. * p < 0.05, ** p < 0.01, *** p < 0.001. Unpaired two-tailed Student’s t-test for (B)-(E). One-way ANOVA followed by Dunnett’s multiple comparisons test for (F), (G).

Then, we investigated the effects of these inhibitors on the amount of cellular ATP. Pretreatment with DCA or oxamate did not affect ATP levels under isoosmotic conditions, suggesting that ATP produced by aerobic glycolysis could be compensated (Fig EV4). In contrast, the relative ATP level upon hyperosmotic stress was significantly decreased by pretreatment with these inhibitors compared with the control (Fig 4F). Considering that the production rate of ATP is approximately 100 times faster in glycolysis than in OXPHOS (Liberti and Locasale, 2016), the result suggests that Warburg-like remodeling upon hyperosmotic stress is induced to compensate for ATP demand under stress conditions by rapidly producing ATP in glycolysis (Leslie *et al*, 2019).

Considering that there was almost no difference in the amount of ATP of control cells under hyperosmotic conditions compared with isoosmotic conditions (Fig 4F, EV4), it is suggested that ATP is consumed as rapidly as ATP is produced by aerobic glycolysis under hyperosmotic stress. As hyperosmotic stress causes DNA damage (Burg, Ferraris and Dmitrieva, 2007) and ATP is required for DNA repair (Lans, Marteijn and Vermeulen, 2012), it is possible that ATP might be consumed for the repair of DNA damage upon hyperosmotic stress.

Finally, we examined the impact of the inhibition of metabolic remodeling on cell death under hyperosmotic conditions. Cell death evaluated by PI staining was not detectable under isoosmotic conditions even in the presence of DCA or oxamate (Fig 4G). However, PI-positive cells were significantly increased by pretreatment with DCA or oxamate under hyperosmotic stress (Fig 4G, H). These data suggest that activation of aerobic glycolysis is necessary for maintaining the ATP level and cell survival under hyperosmotic conditions.

In summary, our findings revealed rapid metabolic remodeling upon osmotic stress through PDK-dependent PDH phosphorylation. Although there have been reports suggesting that hyperosmotic stress enhances aerobic glycolysis (Amara, Zheng and Tiriveedhi, 2016) and suppresses mitochondrial respiration (Grauso *et al*, 2019; Geisberger *et al*, 2021) after several hours or days, we firstly revealed in this study using cancer and macrophage cell lines that cells immediately respond to osmotic stress in a few minutes via PDH phosphorylation and trigger glucose metabolic remodeling. Similar to the general Warburg effect (Nurbubu, 2020), the activation of aerobic glycolysis under hyperosmotic conditions may be derived not only from PDH suppression but also from the activation of enzymes such as PFK, as we mentioned in Fig 2. However, the activation mechanism of these enzymes in hyperosmotic stress may be different, because the response is too immediate to undergo transcriptional changes seen in the general Warburg effect (Hitosugi and Chen, 2014).

Elucidation of the acute regulatory mechanism of PDK in mitochondria will provide a novel methodology to control cellular metabolism without altering osmolarity. Further studies on where and which cells exhibit osmotic stress-dependent metabolic remodeling in vivo will provide new insights into the rapid regulation of cellular metabolism by osmolarity, which might contribute to the treatment of cancer, immune diseases, and other metabolic diseases.

## Materials and Methods

### Cell culture

HeLa cells were cultured in DMEM-low glucose (Sigma, Cat#D6046 or Wako, Cat#041-29775) supplemented with 10% fetal bovine serum (FBS). RAW264.7 cells were cultured in RPMI-1640 medium (Sigma, Cat#R8758 or Wako, Cat#189-02025) supplemented with 10% FBS. All cells were cultured in 5% CO_2_ at 37°C.

### Osmotic stress treatments

Osmolality of the medium is approximately 300 mOsm. 2.5 M NaCl (Wako, Cat#195-01663) solution was added to the medium at final concentrations of 25 mM, 50 mM, and 100 mM, resulting in hyperosmotic stress of approximately 350 mOsm, 400 mOsm, and 500 mOsm, respectively. By adding 2.5 M sorbitol (Wako; Cat#198-03755) solution to the medium at a final concentration of 200 mM, a hyperosmotic stress of approximately 500 mOsm was applied. Hypoosmotic stress was applied by adding distilled water (MilliQ) in the medium to the osmolality of 200 mOsm, 225 mOsm and 250 mOsm. Absolute osmolality was verified by Osmomat 030 (Gonotec).

### Antibodies

The following antibodies were used in this study: Rabbit monoclonal anti-phospho-PDH-E1α (Ser293) antibody (EPR12200: Abcam, Cat#ab177461), mouse monoclonal anti-PDH-E1α antibody (D-6: Santa Cruz Biotechnology, Cat#sc-377092), rat monoclonal anti-α-tubulin antibody (YL1/2: Santa Cruz Biotechnology, Cat#sc-53029), mouse monoclonal anti-actin antibody (AC-40: Sigma, Cat#A3853), mouse monoclonal anti-p38α MAPK antibody (L53F8: Cell Signaling, Cat#9228), rabbit polyclonal anti-COX? antibody (proteintech, Cat#11242-1-AP), mouse monoclonal anti-phospho-p44/42 MAPK (Erk1/2) (Thr202/Tyr404) antibody (E10: Cell signaling, Cat#9106), rabbit polyclonal anti-p44/42 MAPK (Erk1/2) antibody (Cell Signaling, Cat#9102), rabbit polyclonal anti-phospho-acetyl-CoA-carboxylase (Ser79) antibody (Cell Signaling, Cat#3661), rabbit monoclonal anti-acetyl-CoA-carboxylase antibody (C83B10: Cell Signaling, Cat#3676), rabbit polyclonal anti-phospho-Akt (Ser473) antibody (Cell Signaling, Cat#9271), rabbit monoclonal anti-Akt antibody (11E7: Cell Signaling, Cat#4685), and rabbit polyclonal anti-phospho-4E-BP1 (Ser65) antibody (Cell Signaling, Cat#9451). Goat anti-rabbit IgG, HRP-linked antibody (Cell Signaling, Cat#7074), horse anti-mouse IgG, HRP-linked antibody (Cell Signaling, Cat#7076), and goat anti-rat IgG, HRP-linked antibody (Cell Signaling, Cat#7077) were used as secondary antibodies.

### Measurement of oxygen consumption rate (OCR) and extracellular acidification rate (ECAR)

OCR and ECAR measurements were performed using the XF24 or XF96 Extracellular Flux analyzer (Seahorse Bioscience). Cells were plated into XF24 (Agilent; Cat#100867-100) or XF96 (Agilent; Cat#102601-100) cell culture plates and cultured for 24 hr. Prior to performing the assay, culture medium in the wells was exchanged with the Seahorse XF DMEM medium (Agilent, Cat#103575-100) containing 1 g/L glucose (Wako, Cat#041-00595), 2 mM glutamine (Sigma, Cat#G9273), and 1 mM sodium pyruvate (Sigma, Cat#P2256) for HeLa cells or Seahorse XF RPMI medium (Agilent, Cat#103576-100) containing 2 g/L glucose and 1 mM glutamine for RAW264.7 cells. While sensor cartridges were calibrated, cell plates were incubated in a 37°C, CO_2_ free incubator for 60 min prior to the start of the assay. All experiments were performed at 37°C. Each measurement cycle consisted of a mixing time of 3 min, a waiting time of 2 min, and a data acquisition time of 3 min. Time points of OCR and ECAR data refer to the average rates during the measurements cycle. All compounds were prepared at appropriate concentrations in desired assay medium. In a typical experiment, 3 baseline measurements were taken prior to the addition of any compound, and 3 response measurements were taken after the addition of each compound. The OCR and ECAR data were analyzed using Wave software (Agilent).

For the experiment of measuring maximal respiration dependent on fatty acid metabolism (Fig EV2C), HeLa cells were seeded in XF96 cell culture plate. 1 day after seeding, culture medium in the wells was exchanged with DMEM medium (Sigma, Cat#D5030) containing 0.5 mM glucose, 1 mM glutamine, 0.5 mM carnitine (Sigma, Cat#C0283), 1% FBS, 15 mg/L phenol red (Wako, Cat#165-01121), and 3.7 g/L NaHCO_3_ (Nacalai Tesque, Cat#312-13). Next day, prior to performing the assay, the medium was exchanged with assay medium (111 mM NaCl, 4.7 mM KCl, 1.25 mM CaCl_2_, 2.0 mM MgSO_4_, 1.2 mM NaH_2_PO_4_, 2.5 mM glucose, 0.5 mM carnitine, and 5 mM HEPES). 150 µM palmitate-BSA (Agilent, Cat#102720-100) was added just before starting the assay. 40 µM (+)-etomoxir (sodium salt) (Cayman Chemical, Cat#11969), if present, was added 15 min before starting the assay.

The reagents pretreated 30 min before the assay include: 50 mM sodium dichloroacetate (Sigma, Cat#347795), 50 mM sodium oxamate (Wako, Cat#327-24621). The reagents treated for some indicated experiments include: 3-5 µM oligomycin A (Sigma, Cat# 75351), 1 µM (for HeLa cells) or 0.6 µM (for RAW264.7 cells) carbonyl cyanide *p*-trifluoromethoxyphenylhydrazone (FCCP) (Cayman, Cat#15218), 2 µM rotenone (Sigma, Cat#R8875), 4 µM antimycin A (Sigma, Cat#A8674), 30 mM 2-deoxy-glucose (Tokyo Kasei, Cat#D0051), 20 µM (+)-etomoxir (sodium salt). Reagents were injected from the reagent ports automatically to the wells at the time as indicated in the Figures.

### Metabolome analysis using a fully labeled ^13^C_6_-glucose tracer

HeLa cells were seeded at 7×10^5^ cells on a 10 cm dish and cultured overnight. Three dishes (triplicates) were prepared for each condition. Next day, culture media were replaced with 8 ml of ^13^C_6_ glucose-containing culture media. Osmotic stress was started at the same time by adding 0.5 ml of 150 mM (for 300 mOsm) and 1900 mM (for 500 mOsm) sodium chloride solution for isoosmotic stress and hyperosmotic stress, respectively. After 10, 30 and 60 min of osmotic stress, cells were washed twice with 5% mannitol solution and lysed in 1 mL methanol containing 25 μM internal standards (L-methionine sulfone, 2-(N-morpholino) ethanesulfonic acid and D-camphor-10-sulfonic acid). After incubating for 10 minutes at room temperature, the cells and supernatants were collected and stored at -80°C until analysis.

Charged metabolites were extracted from 400 μL lysate with 400 μL of chloroform and 200 μL of Milli-Q water, passing 400 μL of the aqueous phase through a 5 kDa-cutoff spin-filter column. The filtrate was dried using an evacuated centrifuge and resuspended in 25 μL Milli-Q water containing 200 μM reference compounds (3-aminopyrrolidine and Trimesate) prior to analysis by MS.

All the charged metabolites in samples were measured by CE-TOFMS (Agilent Technologies). Detailed conditions for CE-TOFMS-based metabolome analysis were described previously (Satoh *et al*, 2017). For CE-TOFMS system control and data acquisition, we used our proprietary software MasterHands.

For data analysis, the amount of each metabolite was normalized by the average of the total amount at 10 min under isoosmotic conditions. Then, the triplicates in each stimulation condition were averaged and compared. Metabolites exhibiting the coefficient of variation (CV) of triplicate less than 0.4 in all six conditions (3 time points in iso- and hyper-osmotic conditions each) were used for statistics. Graphs were drawn by using R software.

### Nontargeted lipidome analysis

HeLa cells were seeded at 1.4×10^6^ cells on a 10 cm dish. Three dishes (triplicates) were prepared for each condition (200 mOsm hypoosmotic, 300 mOsm isoosmotic, and 500 mOsm hyperosmotic stimulation). The background control dishes contained only medium. After 2 days, the dishes were washed twice with PBS, and then replaced with each osmotic buffer adjusted by mannitol. Isoosmotic buffer (300 mOsm) contained 130 mM NaCl, 2 mM KCl, 1 mM KH_2_PO_4_, 2 mM CaCl_2_, 2 mM MgCl_2_, 10 mM HEPES, 10 mM Glucose, and 20 mM mannitol. Hyperosmotic buffer (500 mOsm) contained additional 200 mM mannitol compared with isoosmotic buffer. In hypoosmotic buffer (200 mOsm), mannitol was excluded and NaCl was reduced to 90 mM from isoosmotic buffer. The dishes were incubated at room temperature for 15 minutes and the buffer was aspired. 6 mL of cold MeOH was added to each dish at 4°C. After incubating for 10 minutes at 4°C, the supernatants were collected and stored at -80°C until analysis.

The supernatants were transferred to glass jacket tubes (FCR&Bio). Then, they were concentrated by centrifugation (Labconco) overnight at 4°C. 100 μL of chloroform was added to dried cells, followed by 30 sec sonication. After 60 min incubation at room temperature, 200 μL of methanol containing internal standards was added and vortexed for 10 sec. After 60 min of incubation, 20 μL of Milli-Q water was added, vortexed for 10 sec, and incubated for 10 min at room temperature. The tubes were then centrifuged at 2,000 g for 10 min at 20°C, and 200 μL of supernatant was transferred to LC-MS vials. The final composition ratio of the lipid extracts was chloroform: methanol: water (5:10:1, v/v/v), and the final concentrations of internal standards were prepared to 25 μM for fatty acids (16:0-d3, 18:0-d3) and 5 μM for acyl-carnitine (18:0-d3).

Identification and quantification of lipid related molecules were performed using LC-MS/MS as described previously with some modifications (Imanishi *et al*, 2019; Tsugawa *et al*, 2020). Briefly, 1 μL of lipid extracts was separated by an ACQUITY UPLC BEH C18 column (2.1×50 mm, particle size 1.7 μm) at a flow rate of 300 μL/min at 45°C using an ACQUITY UPLC system (Waters). Solvent A consisted of acetonitrile/methanol/water (20:20:60, v/v/v) and solvent B was isopropanol, both containing 5 mM ammonium acetate and 10 nM EDTA. The solvent composition started at 100% (A) for the first 1 min and was changed linearly to 64% (B) at 7.5 min, where it was held for 4.5 min. The gradient was increased linearly to 82.5% (B) at 12.5 min, followed by 85% (B) at 19 min, 95% (B) at 20 min, 100% (A) at 20.1 min and 100% (A) at 25 min.

Analysis of lipid related molecules was performed using a Triple TOF 6600 system (AB SCIEX) in the negative and positive ion mode with a scan range of *m/z* 70-1,250. Raw data files from the TOF-MS were converted to MGF files using the program AB SCIEX converter for subsequent quantitative analysis with 2DICAL (Mitsui Knowledge Industry). Identification of molecular species was accomplished by comparison with retention times and MS/MS spectra with commercially available standards or reference samples.

For data analysis, the amount of each lipid related molecule was normalized by the average of isoosmotic conditions. Then, the triplicates in each osmotic condition were averaged and compared.

### Live-cell imaging and measurement of FAOBlue fluorescence

For live-cell imaging experiment, HeLa cells were seeded in φ35 mm glass bottom dishes (Matsunami, Cat#D11130H) which were coated with 1% gelatin (Nacalai Tesque, Cat#16605-42) in PBS in advance. 2 days later, the culture medium was aspired and washed with FBS (-) culture medium. Then 1 mL of culture medium containing 20 µM FAOBlue (Funakoshi, Cat#FDV-0033) was added per dish and osmotic stimulation was simultaneously applied. 40 µM etomoxir, if present, was pretreated 60 min before osmotic stimulation. 90 min after osmotic stimulation, the dish was imaged by a TCS SP5 (Leica) confocal-laser scanning microscope equipped with a stage top incubator (Tokai Hit). The cells were observed in 5% CO_2_ at 37°C using an HC PL APO 63 ×/1.40 oil objective (Leica). FAOBlue was excited at 405 nm. Representative images were adjusted linearly to appropriate brightness using Image J software.

For measuring the fluorescence intensity of FAOBlue, HeLa cells were seeded in a 96 well black glass bottom plate (Matsunami, Cat#GP96000) which were coated with 1% gelatin in PBS in advance. 2 days later, the culture medium was aspired and washed with FBS (-) culture medium. Then 100 µL of culture medium containing 20 µM FAOBlue was added per well and osmotic stimulation was simultaneously applied. 40 µM etomoxir, if present, was pretreated 30 min before osmotic stimulation. Osmotic stress was treated for 15, 30, 60, and 90 min. The fluorescence of FAOBlue was measured at the wavelength (Ex/Em = 405/460 nm) using Varioskan Flash (Thermo Fisher Scientific).

### Immunoblotting

Cells or isolated mitochondria were lysed in lysis buffer (20 mM Tris-HCl pH 7.5, 150 mM NaCl, 10 mM EDTA pH 8.0, 1% sodium deoxycholate, 1% Triton X-100, 1 mM phenylmethylsulfonyl fluoride (PMSF), and 5 µg/mL leupeptin) for 20 min at 4°C. The extracts were clarified by centrifugation for 10 min, and the supernatants were sampled by adding an equal volume of 2 × SDS sample buffer (80 mM Tris-HCl pH 8.8, 80 μg/mL bromophenol blue, 28.8% glycerol, 4 % sodium dodecyl sulfate (SDS), and 10 mM dithiothreitol). After boiling at 98°C for 3 min, the samples were resolved by SDS-PAGE and electroblotted onto a FluoroTrans W membrane (Pall, Cat#BSP0161) or an Immobilon-P membrane (Millipore, Cat#IPVH00010). The membranes were blocked with 5% skim milk (Megmilk Snow Brand) in TBS-T (50 mM Tris-HCl pH 8.0, 150 mM NaCl, and 0.05% Tween 20) and then probed with the appropriate primary antibodies diluted by 1st antibody-dilution buffer (TBS-T supplemented with 5% BSA (Iwai Chemicals, Cat#A001) and 0.1% NaN_3_ (Nacalai Tesque, Cat#312-33)). After replacing and probing the appropriate secondary HRP-conjugated antibodies diluted in 5 % skim milk in TBS-T, antibody-antigen complexes were detected by FUSION SOLO S (VILBER) using an ECL system (GE Healthcare). Quantification was performed via densitometry using Fusion Capt software (VILBER). Representative images were adjusted linearly to the appropriate brightness and contrast using Fusion Capt software (VILBER).

### Mitochondria isolation

HeLa cells were collected from a confluent 15 cm dish and centrifuged. The supernatant was discarded, and the precipitated cells were suspended with Buffer A (270 mM Mannitol, 10 mM HEPES/K^+^ pH 7.4, 0.2 mM EDTA/K^+^ pH 8.0, and 0.1% BSA). The centrifugation and resuspension procedures were repeated at least twice to completely replace the culture medium with Buffer A. Then, the cells were disrupted using a homogenizer and centrifuged at 800 g for 10 min at 4°C. The supernatant was decanted into another tube, centrifuged at 3,000 g for 8 min at 4°C, followed by continuous centrifugation at 7,600 g for 5 min at 4°C. After discarding the supernatant, 10 mL of Buffer B (270 mM Mannitol, 10 mM HEPES/K^+^ pH 7.4, and 0.1% BSA) was added, and the precipitates were suspended and centrifuged at 800 g for 10 min at 4°C. The supernatant was decanted into another tube, and centrifuged at 6,700 g for 10 min at 4°C. Finally, 2.8 mL of Buffer B was added to the precipitate (isolated mitochondria) and suspended. Of note, the experimental instruments in the isolation were immersed in Buffer A beforehand to remove Ca^2+^, and all procedures were performed on ice.

After centrifugating the mitochondrial suspension and discarding the supernatant, osmotic buffer was treated to the precipitate and suspended. 200 mOsm osmotic buffer contained 90 mM KCl, 10 mM HEPES/K^+^ pH 7.4, 0.2 mM EDTA/K^+^ pH 8.0, and 0.1% BSA. 300 mOsm, 350 mOsm, and 400 mOsm osmotic buffer contained additional 50 mM, 75 mM, 100 mM KCl respectively compared with 200 mOsm osmotic buffer.

### Measurement of intracellular ATP amount

Intracellular ATP amount was measured using “ Cell” ATP Assay reagent Ver.2 (Wako, Cat#381-09306). HeLa cells were seeded in a 96 well white transparent bottom plate (Greiner Bio-One Cat#655098 or CORNING, Cat#3610) and cultured for 24 hr. 25 mM NaCl (control), 25 mM DCA or 25 mM oxamate was pretreated 60 min before osmotic stress. Then iso-or hyper-osmotic stress was applied for 15, 30, 60, and 120 min. After osmotic stimulation, all medium was removed and 50 µL per well of ATP reaction mixture (culture medium and assay reagent mixed in equal volumes) was added. Samples were incubated at room temperature for 10 min protected from light, and the luminescence was measured using Varioskan Flash. ATP amount at the timepoint of 0 min under isoosmotic conditions without inhibitors was normalized as 1.

### PI staining

HeLa cells were seeded in a 96 well plate and cultured for 24 hr. 25 mM NaCl (control), 25 mM DCA or 25 mM oxamate was pretreated 60 min before osmotic stress. Then iso-or hyper-osmotic stress was applied for another 24 hr. After osmotic stimulation, the cells were incubated with culture medium containing Hoechst 33342 (DOJINDO, Cat#346-07951) at a concentration of 1 µg/mL and propidium iodide (PI) (DOJINDO, Cat#343-07461) at a concentration of 0.3 µ g/mL for 30 min in 5% CO_2_ at 37°C. The plate was measured and analyzed using the ArrayScan VTI with the optimized Cell Health Profiling BioApplication. Briefly, 2 image sets (Channel 1: nuclei stained with Hoechst 33342, Channel 2: PI) were acquired from 9 fields per well. Cells were subsequently identified as targets from Channel 1, and a PI intensity was assigned to each target from Channel 2. Finally, cell death was calculated as the ratio of the number of PI-positive cells to the number of Hoechst-positive cells.

### Statistical analysis

All data are represented as the mean ± SEM. Statistical tests was performed using Microsoft Excel or R software with RStudio. Unpaired two-tailed Student’s t-test (for two groups) and one-way ANOVA followed by Dunnett’s multiple comparisons test (for more than two groups) were used in this study. For all statistical analyses, *p < 0.05, **p < 0.01, ***p < 0.001. p < 0.05 was considered statistically significant.

## Acknowledgements

We thank S. Torii, K. Ohshima, and S. Shimizu (Department of Pathological Cell Biology, Medical Research Institute, Tokyo Medical and Dental University) for teaching us the mitochondria isolation technique, S. Hasegawa and R. Inagi (Division of CKD Pathophysiology, The University of Tokyo) for allowing us to use the equipment of extracellular flux analyzer, and K. Saito, K. Kato, and K. Umetsu (Institute for Advanced Biosciences, Keio University) for performing MS analysis and analyzing data of the metabolome analysis using ^13^C_6_-glucose tracer. We also thank all of the members of Laboratory of Cell Signaling in The University of Tokyo for critical discussions.

This work was supported by the Japan Science and Technology Agency (JST) under the Moonshot R&D-MILLENNIA program (grant number JPMJMS2022-18 to H.I.); by the Japan Agency for Medical Research and Development (AMED) under the Project for Elucidating and Controlling Mechanisms of Aging and Longevity (grant number JP21gm5010001 to H.I.); by the Japan Society for the Promotion of Science (JSPS) under the Grant-in-Aid for Scientific Research on Innovative Areas (KAKENHI; grant number JP17H06419 to I.N., and JP15H05897 to M.A.) and the Grant-in-Aid for Scientific Research (KAKENHI; grant numbers JP18H03995, JP21H04760 to H.I., JP18H02569, JP22H02761 to I.N., JP20K07598 to S.K.); by The Nakatomi Foundation to I.N.; by The Naito Grant for the advancement of natural science to I.N; by the Takeda Science Foundation to S.K.; and by the research funds from the Yamagata prefectural government and Tsuruoka city.

## Author contributions

H. I. and I. N. co-conceived of and supervised this project. T. I. designed and performed most of the experiments and analyzed the data. K. I. and M. A. performed nontargeted lipidome analysis. S. K. and T. S. performed metabolome analysis using ^13^C_6_-glucose. T. I., H. I., and I. N. drafted the manuscript.

## Expanded View Figure legends

**Figure EV1.**
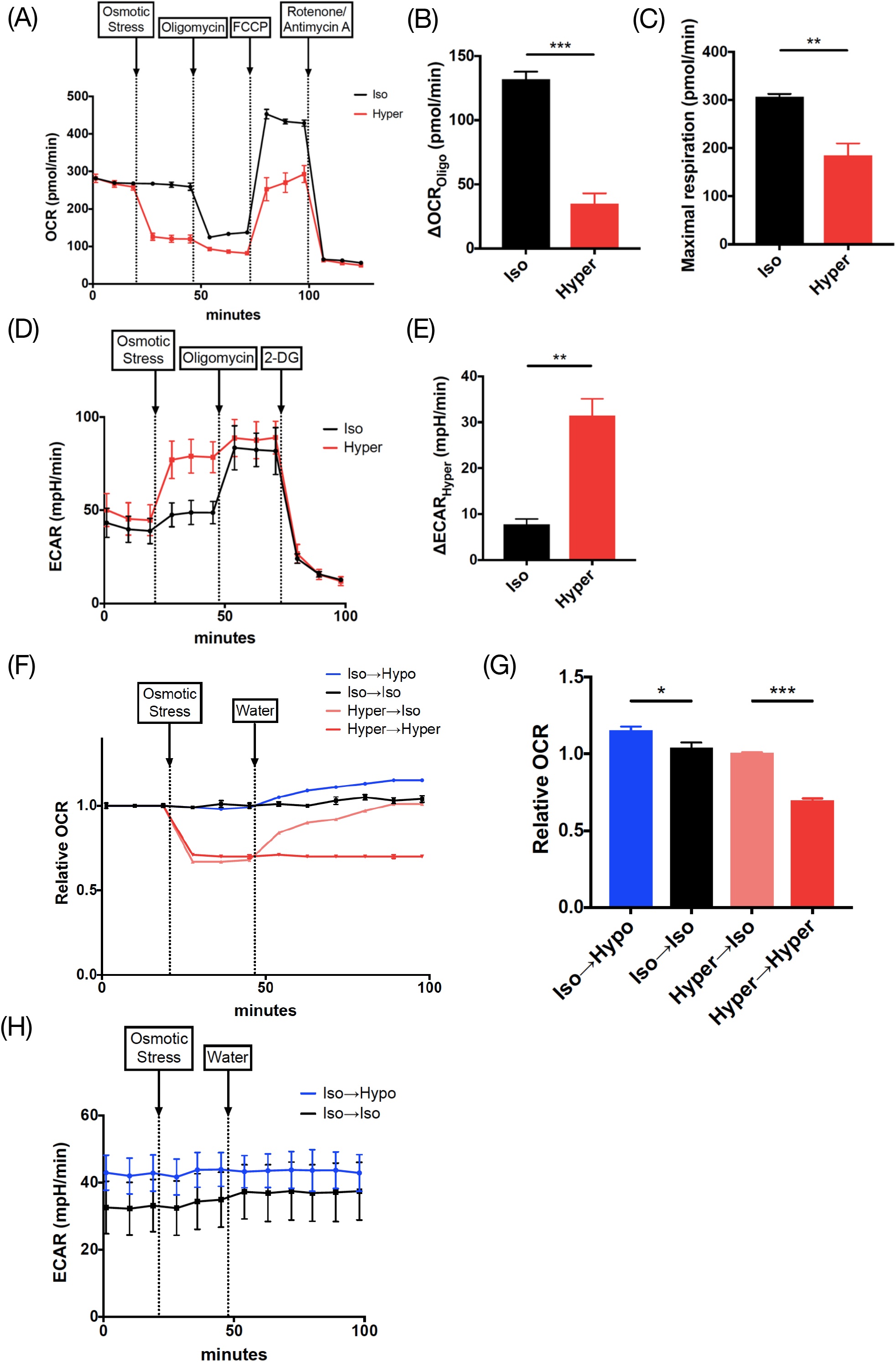
(A) OCR of HeLa cells was measured with sequential treatment with osmotic stress (by adding sorbitol), oligomycin, FCCP, and rotenone/antimycin A. A representative result from three independent experiments is shown. The trace was the average of 4-10 independent wells for each condition. (B) ΔOCR_Oligo_ upon osmotic stress extracted from (A), which was calculated by subtraction between the average of three sequential measurements before and after oligomycin. (C) Maximal respiration upon osmotic stress extracted from (A), which was calculated by subtraction between the average of three sequential measurements before and after FCCP. (D) ECAR of HeLa cells was measured with sequential treatment with osmotic stress (by adding sorbitol), oligomycin, and 2-DG. A representative result from three independent experiments is shown. The trace was the average of 3-5 independent wells for each condition. (E) ΔECAR_Hyper_ extracted from (D), which was calculated by subtraction between the average of three sequential measurements before and after osmotic stress. (F) OCR and (H) ECAR of HeLa cells were measured with the sequential treatment described below for each condition. Iso → Hypo: Medium → Water, Iso → Iso: Medium → Medium, Hyper → Iso: 900 mOsm medium with NaCl → Water, Hyper → Hyper: 900 mOsm medium with NaCl → 400 mOsm medium. The OCR is normalized by the average of the first three measurements. A representative result from three independent experiments is shown. The trace was the average of 3-5 independent wells for each condition. Hyperosmotic stress: 400 mOsm. (G) Values of relative OCR in the last measurement (98 min) extracted from (F). Hypoosmotic stress: 225 mOsm, Isoosmotic stress: 300 mOsm, Hyperosmotic stress: 500 mOsm except (F) and (G). Data are represented as the mean ± SEM. * p < 0.05, ** p < 0.01, *** p < 0.001. Unpaired two-tailed Student’s t-test.

**Figure EV2.**
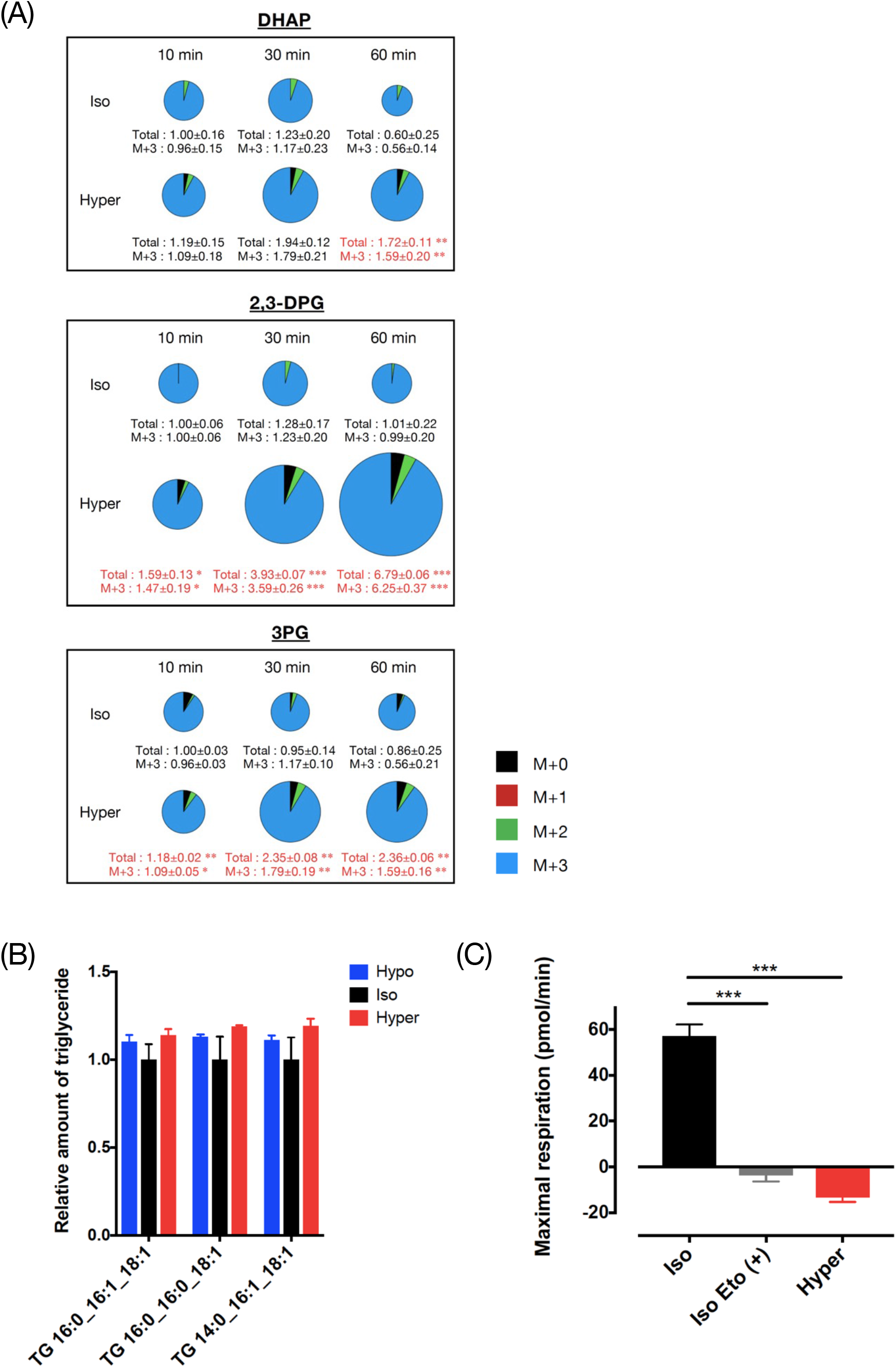
(A) Related to Figure 2A, the relative amounts of metabolites in glycolysis (DHAP, 2,3-DPG and 3PG) are shown. (B) Related to Figure 2D, relative amounts of several triglycerides composed of various combinations of fatty acids in HeLa cells at 15 min after osmotic stress. TG: triglyceride. (C) Maximal respiration using the medium for measuring OCR dependent on fatty acid oxidation (see Methods), which was calculated by subtraction between the average of three sequential measurements before and after FCCP. Three independent experiments were performed in which 5-11 independent wells were measured for each condition. Etomoxir (40 µM) was added 15 min before starting the assay. Palmitate-BSA (150 µM) was added just before starting the assay. Hypoosmotic stress: 200 mOsm, Isoosmotic stress: 300 mOsm, Hyperosmotic stress: 500 mOsm. Data are represented as the mean ± SEM. N=3 (three samples for each condition) except (C). * p < 0.05, ** p < 0.01, *** p < 0.001. Unpaired two-tailed Student’s t-test at the same time points for (A). One-way ANOVA followed by Dunnett’s multiple comparisons test for (B), (C).

**Figure EV3.**
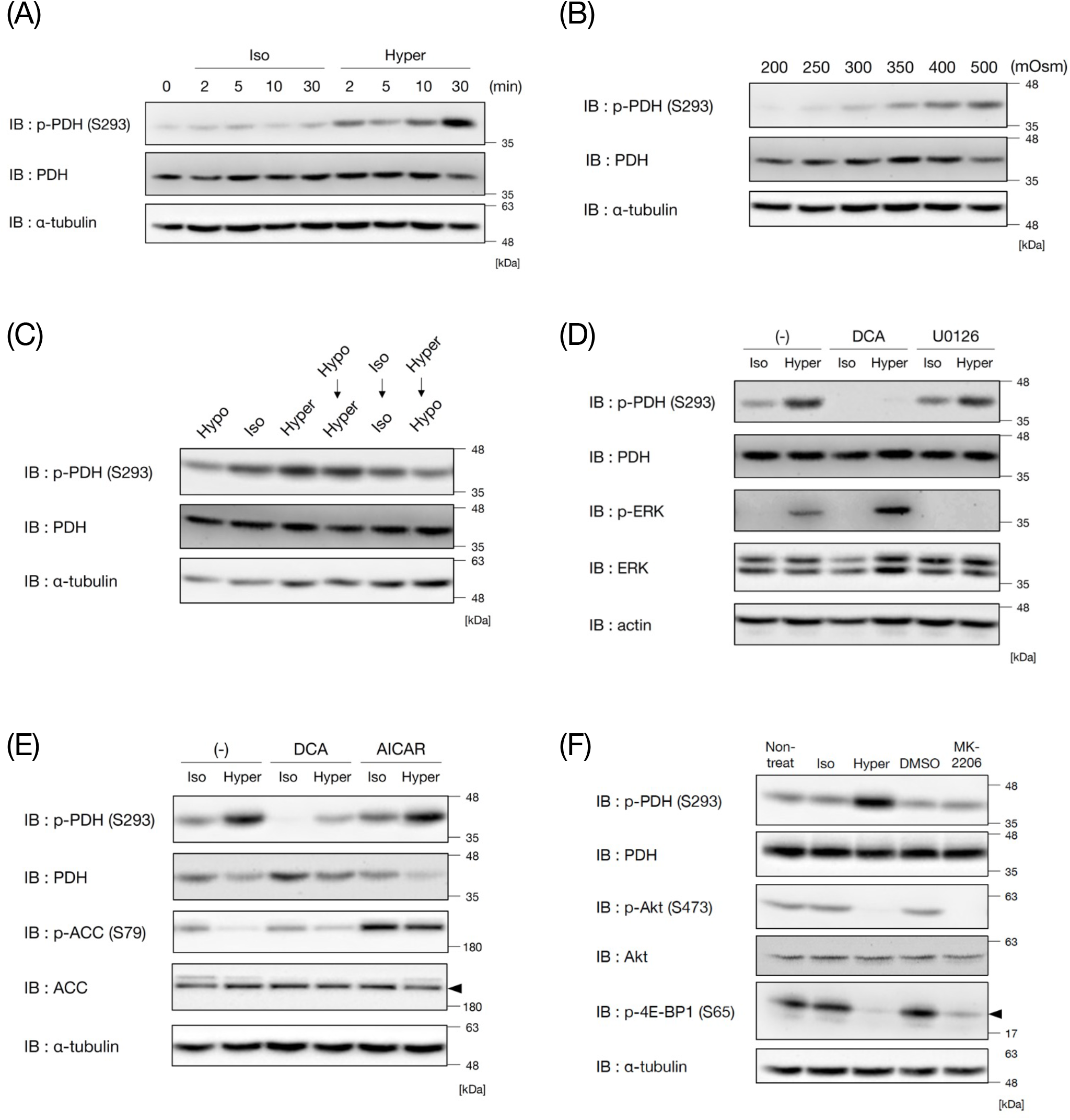
(A) Immunoblot analysis of PDH phosphorylation after 2, 5, 10 and 30 min of osmotic stress by adding sorbitol to HeLa cells. (B) Immunoblot analysis of PDH phosphorylation under various osmolarities (Hypo: 200, 250, Iso: 300, Hyper: 400, 500 mOsm) in HeLa cells. Osmotic stress: 30 min. (C) Immunoblot analysis of PDH phosphorylation in HeLa cells when osmotic stress was changed halfway. Normal osmotic stress for 30 min in the left three lanes (Hypo: 200 mOsm, Iso: 300 mOsm, Hyper: 400 mOsm). In the right three lanes, osmotic stress was altered after 30 min as indicated. (D) Immunoblot analysis of PDH phosphorylation after 10 min of osmotic stress in HeLa cells pretreated with the MEK inhibitor U0126 (20 µM) or DCA (25 mM) for 60 min. (E) Immunoblot analysis of PDH phosphorylation after 30 min of osmotic stress in HeLa cells pretreated with the AMPK activator AICAR (5 mM) or DCA (25 mM) for 60 min. (F) Immunoblot analysis of PDH phosphorylation in HeLa cells after 10 min of osmotic stress and after treatment with the Akt inhibitor MK-2206 (500 nM) for 60 min under isoosmotic conditions. Hypoosmotic stress: 200 mOsm, Isoosmotic stress: 300 mOsm, Hyperosmotic stress: 500 mOsm except (C).

**Figure EV4.**
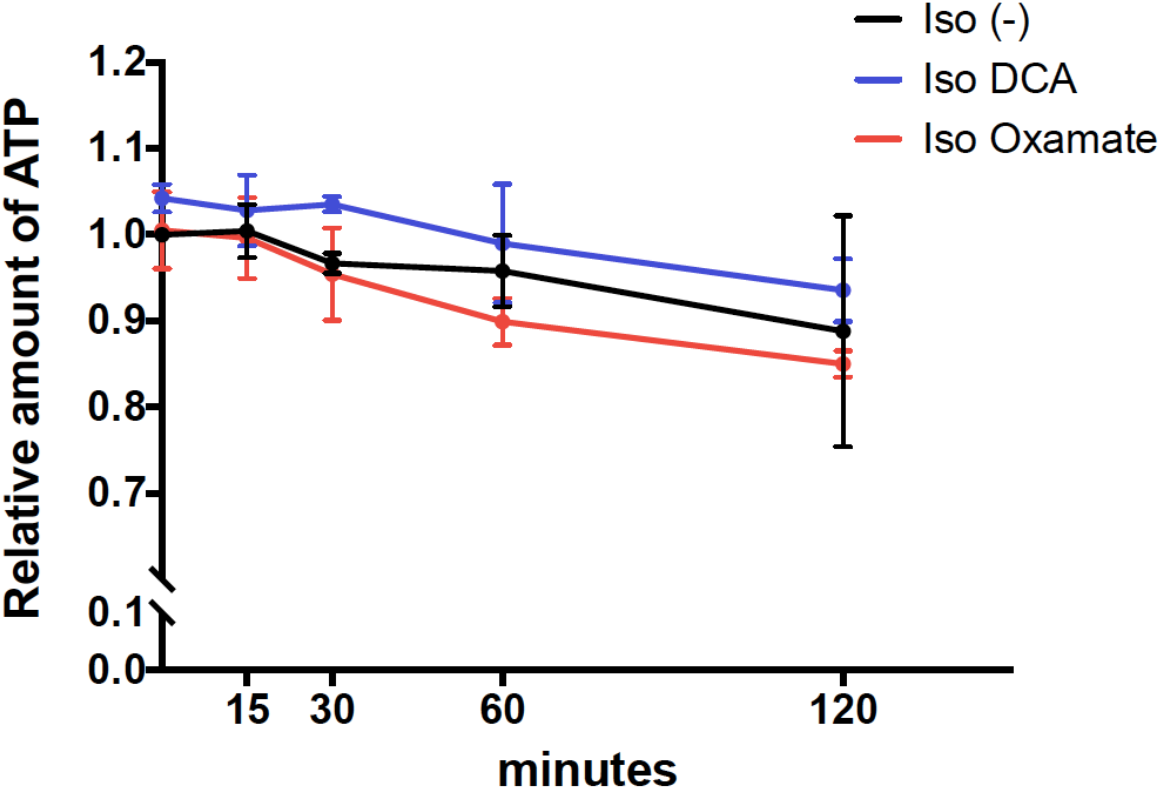
Related to Figure 4F, the relative amount of ATP in HeLa cells was measured under isoosmotic conditions with pretreatment with DCA (25 mM) or oxamate (25 mM) for 60 min using a plate reader. The timepoint of osmotic stimulation was defined as 0 min. Three independent experiments were performed and averaged.

